# Quantum Sensing Reveals Chemotherapy Induced Persistent Oxidative Footprints in Fixed Brain Cancer Cells

**DOI:** 10.1101/2025.07.27.666994

**Authors:** Jyotiprakash Parhi, Arpita Ghosh, Sritama Saha, Disha Zaveri, Ayan Majumder, Siddharth Tallur, Abhijit Majumder, Kasturi Saha

## Abstract

Glioblastoma (GBM) is a highly aggressive cancer with poor prognosis despite temozolomide(TMZ) treatment. TMZ induces oxidative stress, causing persistent redox alterations. We report the first use of nitrogen-vacancy centers in nanodiamonds as quantum sensors for *T*_1_ relaxometry in fixed GBM cells, detecting stable paramagnetic signatures linked to chemotherapy-induced persistent oxidative damage. U87-MG cells treated with TMZ show dose-dependent *T*_1_ shortening, consistent with conventional ROS fluorescence assays. TBHP studies confirm that the *T*_1_ response reflects accumulated redox adducts. Measurements remain robust across fixation and permeabilization protocols. Since drug-resistant GBM cells preserve antioxidant signatures, unchanged *T*_1_ times after treatment may indicate early resistance phenotypes. This fixed-tissue compatible readout could stratify patient response, guide adaptive therapy, and support biopsy-based diagnostics and post-treatment drug analysis in routinely preserved clinical specimens enabling retrospective studies and better outcomes for future personalized oncology decision making pathways.

## Introduction

The emergence of fluorescent nanodiamonds (FNDs) has ushered in a new sensing modality in biomedical engineering, in various cell-based applications in biology, including bio-labeling, drug delivery, and bio-sensing, due to their remarkable photostability and biocompatibility.^1,2^ The substitution of carbon by a nitrogen atom adjacent to a vacancy in the diamond lattice results in the nitrogen vacancy (NV) center, a specific defect in terms of a localized impurity in FNDs, responsible for its unique dynamic opto-magnetic properties. By converting environmental magnetic signals into optical signals, this specific defect attribute enables high-sensitivity detection of nanoscale magnetic fluctuations. The capability of the nitrogen-vacancy (NV) center to detect unpaired electron spins in its environment has been extensively exploited in physics and materials science.^3,4^ It enables the detection and characterization of spin defects, paramagnetic ions, nanostructures, and nanoparticles with high spatial resolution.^5,6^ In particular, *T*_1_ relaxometry—also referred to as longitudinal relaxometry—has emerged as a powerful modality for probing magnetic fluctuations at the nanoscale. As a form of noise spectroscopy,^7^ it provides direct access to the local magnetic noise spectral density, making it highly suitable for investigating dynamic spin environments. This approach has been successfully applied in condensed matter physics to image magnetic domain walls and skyrmions,^8^ and in materials science to probe magnetic noise in superconductors and engineered nanostructures.^9–11^ Furthermore, NV-based *T*_1_ relaxometry has enabled nanoscale magnetic and thermal imaging,^12^ demonstrating its versatility as a quantum sensing platform. Building on these advances, nanodiamonds hosting NV centers (FNDs) extend these capabilities into complex and heterogeneous environments. Their biocompatibility and optical addressability allow their integration into diverse systems while retaining quantum sensing functionality. This positions *T*_1_ relaxometry with FNDs as a uniquely adaptable technique that bridges fundamental physics and applied sensing. In the biological context, this technique has been most extensively explored due to its ability to operate under ambient and physiological conditions. NV-based *T*_1_ relaxometry has been demonstrated as an effective tool for detecting free radicals and paramagnetic species within living cells.^13–17^ Free radicals, characterized by unpaired electrons, are inherently paramagnetic and play critical roles in cellular processes such as aging,^18^ immune response,^19^ and cancer progression.^20^ However, an imbalance in free radical levels can disrupt cellular homeostasis and damage organelles, emphasizing the need for sensitive methods for their detection and quantification. The ability of *T*_1_ relaxometry to sensitively probe their associated magnetic noise provides a non-invasive route to monitor oxidative stress at the nanoscale.^21,22^ Beyond radical detection, fluorescent nanodiamonds have also been employed for intracellular temperature mapping, pH sensing when functionalized with responsive polymers, and tracking particle orientation^23,24^

Currently, the vast majority of quantification in this burgeoning field is conducted in living cells to monitor real-time ROS generation and the immediate radical response to cellular stressors.^15,25–27^ For example, measuring the nanoscale distribution of free radicals generated during bacterial phagocytosis in macrophages or the metabolic fluctuations in primary human dendritic cells.^15,25,26,28^ While these live-cell measurements provide invaluable kinetic insights, they are fundamentally tethered to the transient nature of primary ROS. Short-lived species such as the hydroxyl radical (*·OH*) or superoxide anion (*O^·−^*) have biological half-lives ranging from nanoseconds to microseconds, necessitating detection at the exact moment of generation. However, a critical and often overlooked aspect of redox biology is that ROS not only exist as fleeting intermediates, but also, their activity results in permanent, stable molecular modifications within the cellular architecture. These “oxidative footprints” include irreversible alterations such as protein carbonylation, lipid peroxidation, and the formation of stable DNA adducts like 8-hydroxydeoxyguanosine (8-OHdG). ^29–35^ These modifications persist long after the initiating reactive species have been neutralized or have decayed. Crucially, many of these secondary products exhibit paramagnetic properties or lead to the sequestration of paramagnetic transition metals (such as *Fe*^3+^ or *Cu*^2+^) within the fixed cellular matrix.^36–38^ Consequently, even in fixed cell samples, these stable redox-driven modifications provide a reservoir of magnetic noise that can be captured and detected by FNDs. The ability to detect these permanent, irreversible oxidative signatures in fixed tissues represents a powerful, yet unexplored, modality in quantum sensing that offers compatibility with standard clinical pathology workflows.

Glioblastoma (GBM) is one of the most aggressive forms of brain cancer, known for its rapid growth and resistance to treatment.^39^ Globally, around 3.19 per 100,000 people are diagnosed each year, with a median survival period of 12-15 months.^40^ Temozolomide (TMZ) is the standard chemotherapy used for GBM, often administered in conjunction with radiation therapy.^41^ Although it can slightly extend survival, many patients develop resistance, limiting its long-term effectiveness. TMZ works by adding methyl groups to DNA, particularly at the O6 position of guanine, leading to DNA damage and cell death.^42^ This alkylation triggers oxidative stress and increases reactive oxygen species (ROS) within tumor cells.^42^ Elevated ROS contributes to cytotoxicity, but can also activate survival pathways, complicating treatment outcomes. In parent GBM cells, TMZ-induced ROS levels typically surge, contributing to DNA damage and cell death. However, in relapsed or resistant cells, ROS levels often decrease due to enhanced antioxidant defenses, allowing cells to survive and evade the cytotoxic effect of TMZ. This redox shift is a key feature of treatment resistance.^43,44^ Beyond the initial surge of transient reactive species, the clinical efficacy of TMZ is profoundly linked to the accumulation of stable, redox-driven molecular damage, specifically lipid peroxidation and protein carbonylation, which serve as permanent oxidative footprints within the cell. While primary ROS dissipate rapidly, their reaction with cellular membranes generates reactive aldehydes, such as 4-hydroxynonenal (4-HNE) and malondialdehyde (MDA), as a result of lipid peroxidation. They form stable covalent adducts with the matrix and contribute significantly to TMZ-induced cytotoxicity. These modifications are effectively frozen in place by standard fixation protocols and, crucially, they retain paramagnetic properties that generate a measurable reservoir of magnetic noise even after the ROS is gone. In this work, we show that FNDs can uniquely step in by utilizing *T*_1_ relaxometry in fixed GBM cells. We demonstrate that NV centers can capture these persistent oxidative signatures that traditional fluorescent probes (like DCF) cannot accurately detect post-fixation. Thus, quantum sensing approach allows for the quantification of drug-induced damage, providing a tissue-compatible readout to stratify patient response and identify early resistance phenotypes where the oxidative footprint is absent due to enhanced antioxidant defenses (Fig.1a).

**Figure 1:**
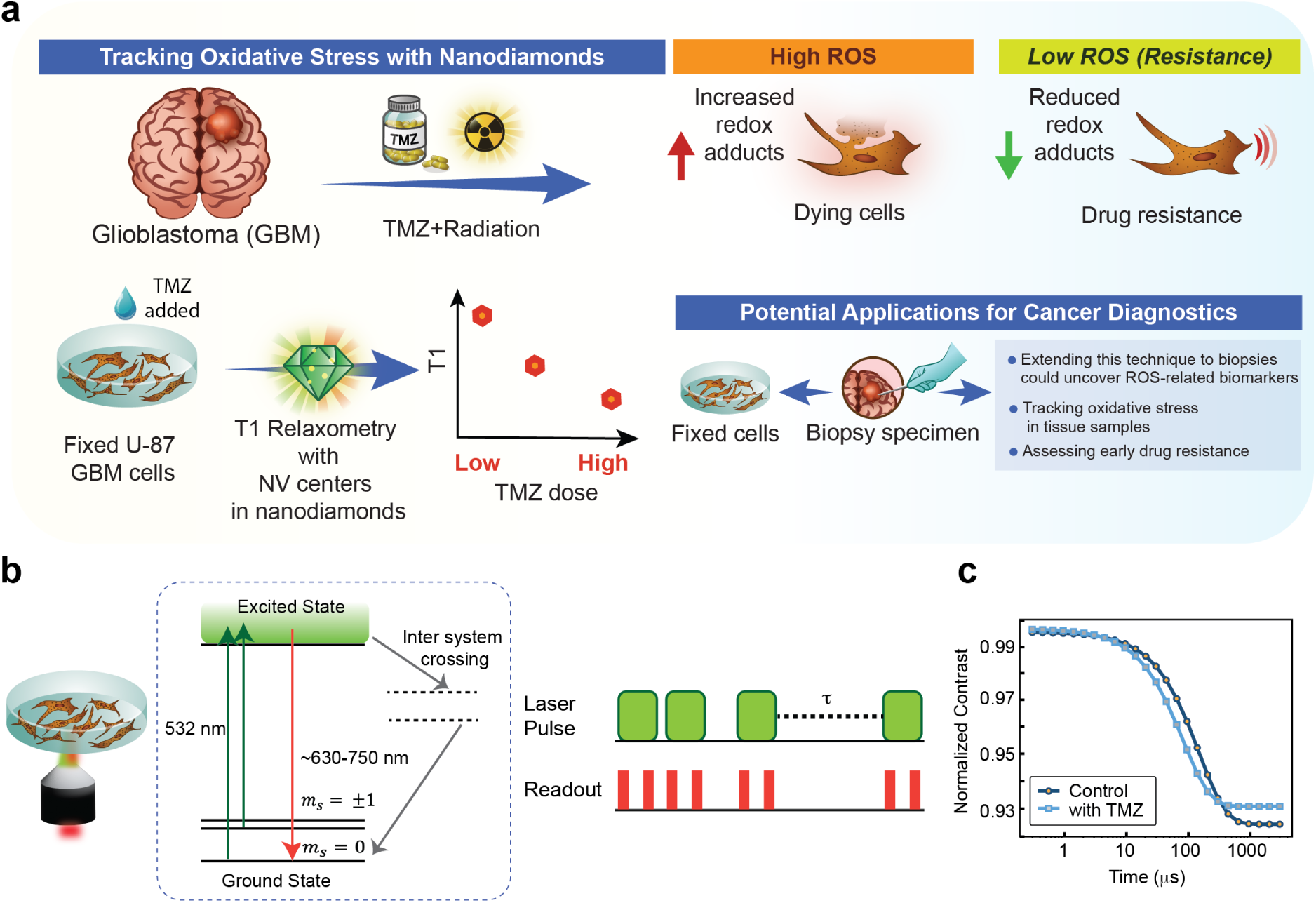
Relaxometry and NV energy levels along with TMZ-induced ROS-mediated redox adducts generation in GBM cells. (a) Schematic illustration of ROS-mediated redox adducts generation induced by TMZ in GBM cells and its effect on local magnetic noise, as probed by nanodiamond-based relaxometry, highlighting potential applications. (b) Schematic of the microscope setup(left), showing the objective lens, cell sample, and the green excitation and red emission light paths. Energy level diagram of the NV center(middle), showing optical excitation (green arrows), radiative decay (red arrows), and non-radiative intersystem crossing pathways. The *T*_1_ relaxometry pulse sequence(right) is depicted, where green bars represent laser (532 nm) pulses that optically initialize the NV spin into the bright, polarized state (*m_s_* = 0). During the variable dark interval *τ*, the spin stochastically relaxes toward a thermal mixed state (*m_s_* = 0*, ±*1). Readout pulses (red bars) measure the resulting fluorescence contrast. (c) *T*_1_ relaxometry curves (log scale) comparing control and TMZ-treated conditions. Showing a clear reduction in *T*_1_ following TMZ treatment reflects an increased concentration of local magnetic species.

We first assessed the accurate measurement of intracellular ROS, standardized ND treatments by drop-casting NDs without cells, and optimized specific experimental parameters like readout time, polarization time, and time step for incrementing dark time (Fig.1b). ND uptake was evaluated and quantified *T*_1_ values in cells after 2 hours of ND treatment. MTT assays confirmed the biocompatibility of NDs over 24 hours, and additional experiments were performed to assess ND distribution area, photon count, and retention in paraformaldehyde (PFA)-fixed cells at time points ranging from 8 to 74 hours. To achieve reliable *T*_1_ relaxometry measurements, we also addressed ND aggregation by optimizing the delivery protocol. Then, in the U87-MG GBM cell line, which represents grade IV astrocytoma, we assessed the irreversible oxidative footprints due to ROS following TMZ treatment at concentrations of 250 µM and 1000 µM. We used NDs as intracellular sensors to track the redox adduct levels through *T*_1_ relaxometry in the TMZ-treated cells, compared to untreated control cells (Fig.1c). To validate these findings, we also analyzed the samples using the conventional DCFDA assay to measure the nascent ROS levels. Suitable control experiments were also employed to ensure that the *T*_1_ signals were generated from the ROS-mediated changes in the cells and not cellular artifacts due to fixation or permeabilization methods, or the time of application of NDs (i.e., in live cells or after fixation).

Together, these observations highlight a critical gap in current redox sensing approaches, which predominantly focus on transient ROS and lack the ability to capture persistent oxidative damage in clinically relevant fixed samples. We therefore hypothesize that *T*_1_ relaxometry using FNDs can detect stable, lipid peroxidation–derived oxidative footprints retained in fixed GBM cells following TMZ treatment. For the first time, we establish and optimize an FND-based sensing framework for reliable detection of these persistent redox signatures and evaluate their potential as quantitative indicators of drug-induced damage and early resistance phenotypes. This approach introduces a new paradigm for retrospective redox analysis in fixed biological specimens and offers a promising strategy for integrating quantum sensing into routine pathology-compatible workflows.

## Results

### Cellular Uptake and Cytotoxic Effects of Nanodiamonds in U87-MG GBM Cells

To evaluate whether NDs were internalized by U87-MG cells, we treated cells with both 35 nm and 70 nm size NDs (SIGMA) at concentrations of 2.5, 5, and 10 *µ*g/mL. Following a 2-hour incubation, cells were fixed and imaged (Fig.2a). We confirmed that the NDs were internalized and not merely adhered to the cell surface by performing z-stack imaging. (see Supplementary Fig.S1, data shown for 10 *µ*g/mL, and for 70 nm size NDs). In addition, to assess whether internalized NDs had any cytotoxic effects, MTT assay was performed after 24 hours of ND treatment across a 0–10 *µ*g/mL concentration range for both 35 nm and 70 nm particles (Fig.2d). The MTT results showed no significant change in cell viability at any of the tested concentrations, indicating good cytocompatibility of the NDs up to 24 hours. The possibility that the observed lack of toxicity arose from cellular clearance of the nanodiamonds, rather than their intrinsic biocompatibility, was excluded by examining fixed cells 24 hours after treatment with 10 µg/mL NDs (Fig.2b,c). NDs remained clearly visible inside the cells, confirming their retention over this time period. Nanodiamonds are efficiently internalized within approximately 2 hours, and their continued presence up to 24 hours reflects their intracellular retention rather than any ongoing cellular metabolism. Following fixation, cells no longer exhibit metabolic activity, yet the internalized NDs remain visible because fixation preserves their spatial localization. Thus, the observation of NDs at later time points, including post-fixation, is a consequence of their stable retention within the cellular environment and not a marker of metabolic function. Taken together, these findings suggest that the NDs are efficiently internalized, retained within cells for at least 24 hours, and do not adversely affect cell viability. Although the initial experiments were carried out using both 70 nm and 35 nm FNDs, a more careful review of the literature indicated that the smaller particles tend to emit weaker fluorescence signals, making it difficult to acquire reliable data within a reasonable time frame. Additionally, their increased mobility poses challenges to stable tracking.^15^ Taking these factors into account, along with insights from our preliminary data, we proceeded with the 70 nm NDs for all subsequent studies.

**Figure 2:**
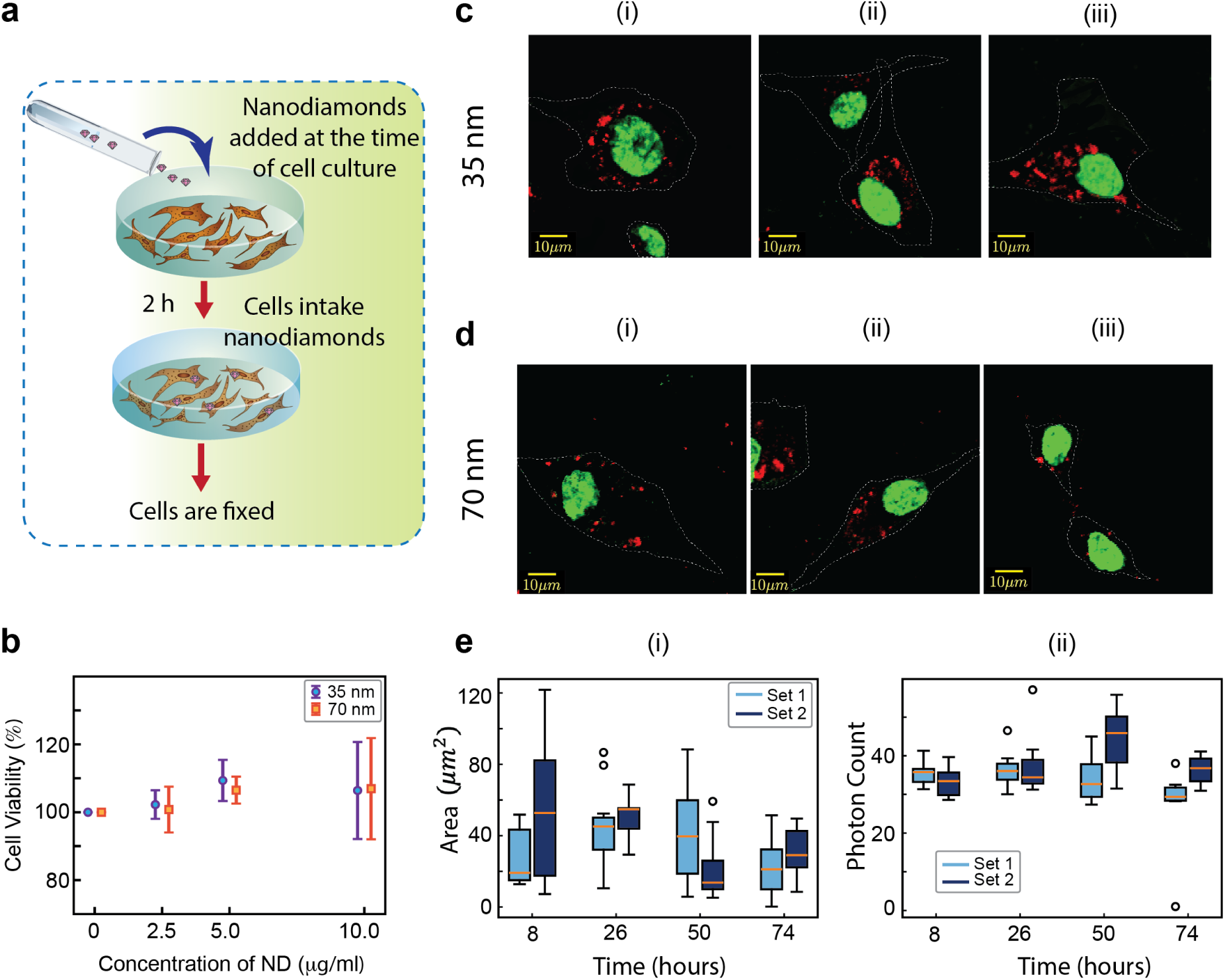
Cellular Uptake and Cytotoxic Effects of Nanodiamonds in U87-MG GBM Cells. (a) Schematic illustration of cellular uptake of NDs. (b) MTT assay for 35nm and 70nm NDs internalized for 24 hours, at concentrations ranging from 0-10 µg/mL, in U87-MG cells to assess cell viability. Error bars represent ± SEM across three independent biological replicates. (c,d) Cellular internalization of 10 µg/mL NDs: 35 nm (c) and 70 nm (d) was assessed 24 hours post-treatment to confirm that the observed cell viability at the highest ND concentration was not due to expulsion of the NDs from the cell. The scale bar corresponds to 10µm. Pseudo colors green denote Hoechst-stained nucleus, and red denotes signal from NDs. The faint white dashed line represents the cell boundary obtained from the brightfield image. (e) (i) Average area occupied by nanodiamonds inside cells as seen after 8, 26, 50, and 74 hours for two replicates. (ii) Average fluorescence (in photon counts) from nanodiamonds as seen after 8, 26, 50, and 74 hours for two replicates.

### Intracellular Characterization of Nanodiamonds in U87-MG Cells for Reliable *T***_1_** Relaxation Measurements

The intracellular behaviour of nanodiamonds over time was investigated by treating U87-MG cells with 70 nm nanodiamonds at a concentration of 10 *µg/mL*. Cells were fixed 2 hours after nanodiamond exposure and subsequently imaged at multiple time points ranging from 8 to 74 hours post-treatment. The area occupied by NDs in cells, as well as photon counts, was examined to assess their retention over time. An overlay of images shows that NDs have been ingested in the cell and cluster near the nucleus. While there is variation in the area occupied by NDs in the cells, there is no trend between area and time of imaging(Fig.2d(ii)). This could be attributed to differences in cell size. The average photon counts also do not show any trend with time and are 28.11 *±* 4.50, 32.73 *±* 5.23 from sets 1 and 2, respectively (Fig.2d(iii)). Thus, NDs are retained in the cells for at least 74 hours after fixation, and neither area nor intensity depends on time, allowing *T*_1_ measurements to be carried out for up to 3 days after fixation. Additionally, we aimed to optimize the size and dispersion of ND clusters inside cells, as larger aggregates positioned near smaller ND puncta could potentially interfere with the signal, overshadowing or distorting the actual *T*_1_ measurements. This could lead to inconsistent and unreliable measurements of ROS levels following different cellular treatments. To address this, we adopted a modified ND preparation protocol (detailed in the Methods section). After sonication and brief vortexing, the ND suspension was allowed to stand undisturbed for 30 minutes, during which time the larger aggregates settled at the bottom. Only the supernatant was then used to treat cells at a concentration of 10 *µg/mL*, resulting in a visible reduction in clumping under bright-field microscopy. To further improve dispersion, we reduced the treatment concentration to 5 *µg/mL* while following the same preparation steps. This yielded an even more uniform distribution of NDs with minimal aggregation inside the cells (Supplementary Fig.S2). Overall, this optimized protocol significantly improved intracellular ND dispersion, enabling more accurate and reproducible *T*_1_ relaxometry measurements in U87-MG cells.

### Monitoring TMZ-induced ROS Mediated Oxidative Footprints

The baseline sensitivity is established by using U87-MG cells that were first treated with 100 µM tert-butyl hydroperoxide (TBHP), a potent inducer of oxidative stress and lipid peroxidation in cells.^45^ TBHP-treated cells exhibited a significant reduction in average *T*_1_ relaxation time compared to untreated controls (Supplementary Fig.S3a), confirming an increase in local magnetic noise and validating the utility of ND sensors for monitoring intracellular ROS-mediated oxidative changes at the single-cell level. We then treated the cells with 0 µM (control), 250 µM, and 1000 µM TMZ as shown in Fig.3a. A clear, dose-dependent reduction in mean *T*_1_ relaxation times was observed with increasing TMZ concentrations (Fig.3b). This decrease suggests an elevation in local spin noise sources, attributed to the treatment-induced detection of radicals within the U87-MG cells.^46–48^

**Figure 3:**
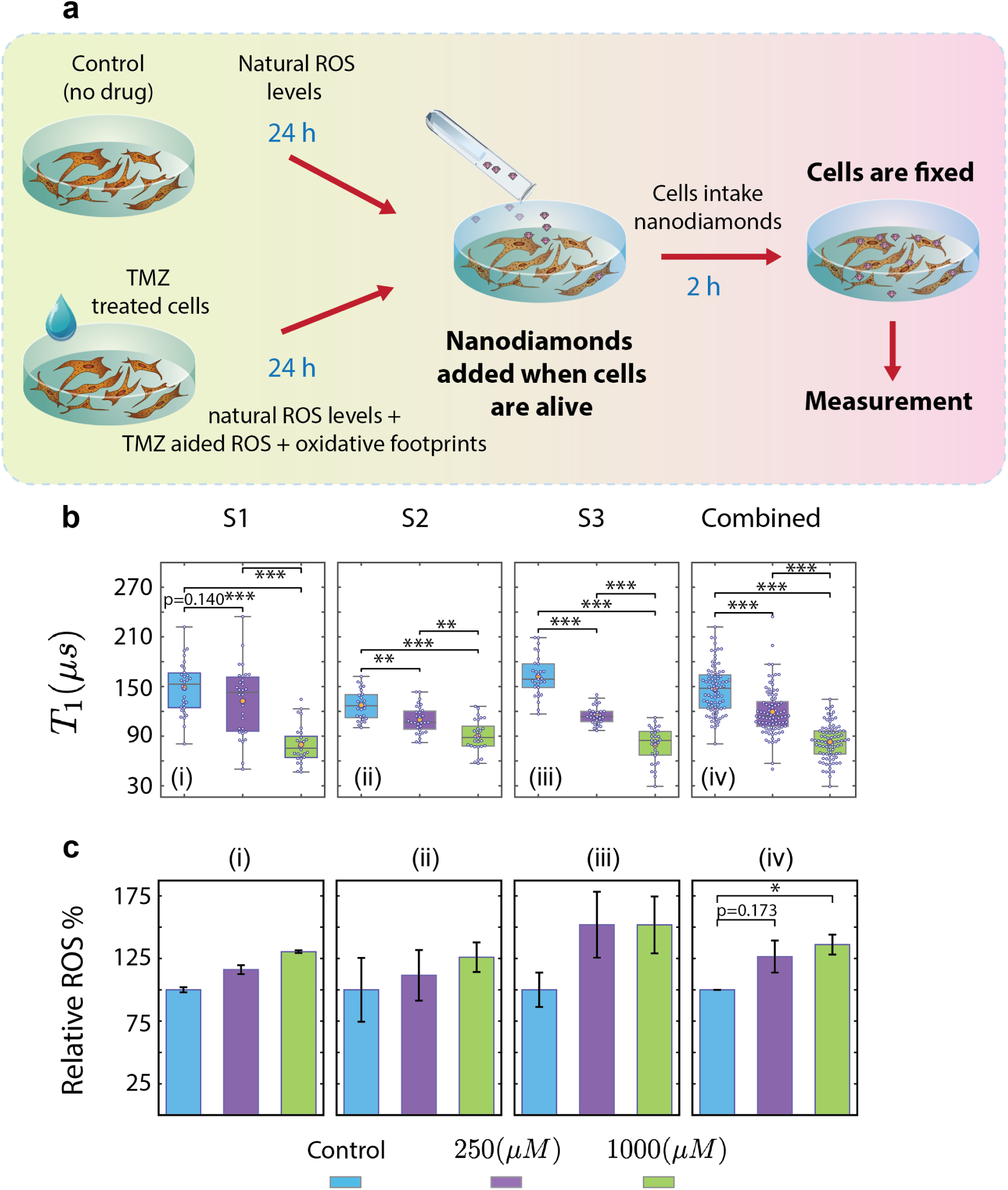
Monitoring TMZ-induced ROS mediated oxidative footprints in GBM cells via ND-based *T*_1_ relaxometry. (a) Schematic illustration of the experimental workflow for sample preparation for relaxometry measurements. (b) *T*_1_ relaxometry measurements for control and TMZ-treated U87-MG cells (250 and 1000 *µ*M) across three independent experimental sets (S1–S3) and the pooled dataset (Combined). Box plots show the distribution of *T*_1_ values (median), with mean (orange dot) and individual measurements (white dots) overlaid. Statistical significance is indicated by adjusted *p*-values (Holm correction); thresholds: *∗ ∗ ∗ <* 0.001, *∗∗ <* 0.01, non-significant comparisons are noted with actual p-values. (c) DCFDA assay to estimate intracellular ROS generation in U87-MG cells under control and TMZ-treated conditions (250 *µ*M and 1000 *µ*M). Data are shown for three biological replicates and for the combined analysis. Error bars represent mean ± SEM *∗ <* 0.05. For both (b) and (c), colors indicate treatment groups as shown in the legend.

We assess the reproducibility of *T*_1_ measurements by performing experiments across three independent biological replicates, capturing at least 30 individual measurements (*∼* 25-27 cells) per condition. While minor variations in the mean *T*_1_ values were observed between sets, this stochasticity is a well-documented characteristic of intracellular NV relaxometry. It arises from the inherent heterogeneity of the biological environment, varying local paramagnetic impurities, and subtle differences in single-particle surface states.^14,15,49,50^ Despite this expected particle-to-particle variation, the overarching dose-dependent reduction in *T*_1_ upon TMZ treatment remained highly robust across all sets. By transparently reporting the full distribution of single-particle data alongside the statistical means and error bars (Fig.3), we demonstrate that NV-based relaxometry effectively and reliably captures biologically significant metabolic shifts at the nanoscale, overcoming challenges posed by single-cell variability. Further, ROS generation was confirmed by performing DCFDA (2’,7’-dichlorodihydrofluorescein diacetate) assay in live cells. TBHP treatment led to a *∼*6-fold increase in ROS compared to control (Supplementary Fig.S3b). For TMZ-treated cells, each of the three biological replicates showed a progressive increase in ROS with increasing drug concentration (Fig.3c(i-iii)). The cumulative data showed a *∼*30% and *∼*40% increase in ROS levels with 250 µM and 1000 µM TMZ treatments, respectively, compared to the untreated control (Fig.3c(iv)), thereby validating the *T*_1_ relaxometry findings as an outcome of the ROS-mediated changes.

### Quantitative and Robust Detection of TMZ-induced Oxidative Stress

A quantitative comparison between NV-based *T*_1_ relaxometry and the conventional DCFDA assay was performed by analysing their normalized responses across TMZ treatment conditions using linear regression (Supplementary Fig.S4a,b). This regression serves as a practical metric to assess proportional scaling between the two modalities. Across all three independent datasets (Supplementary Fig.S4a), as well as in the combined analysis (Supplementary Fig.S4b), the slopes consistently exceeded one, supporting the superior sensitivity of NV relaxometry, since both datasets are normalized to the control; a slope greater than unity directly indicates that *T*_1_ relaxometry provides an amplified response compared to DCFDA for the same dose of TMZ treatment. While goodness-of-fit values showed some variability (*R*^2^ from 0.68 to 1.0)—with the lower *R*^2^ in Set 3 primarily due to a flat response of the DCFDA response—the overall consistency across independent replicates (*R*^2^ = 0.78 for combined data) confirms a robust trend. As the regression is based on only three dose points, it should therefore be interpreted cautiously. These findings establish the NV center-based relaxometry as a more sensitive and quantitative measure of intracellular oxidative stress response than DCFDA.

These observations were confirmed to arise from genuine changes in the intracellular spin environment, rather than from other chemical artefacts, through a series of control analyses. In cell-free media supplemented with TMZ, *T*_1_ measurements showed no significant shift (Supplementary Fig.S4c), indicating that the therapeutic drug does not alter the ND relaxation properties in the absence of cells. To minimize intensity-dependent artifacts, measurements were carefully acquired from regions with comparable photon counts across all conditions. Our analysis showed no systematic dependence of *T*_1_ on count rates across all treatment conditions(Supplementary Fig.S5a,b). Spearman correlation and pvalue heatmaps confirmed that the observed *T*_1_ shifts were independent of photon statistics, ensuring that the sensitivity remains robust across minor variation in ND brightness(Supplementary Fig.S5c,d). Taken together, these controls validate that the measured *T*_1_ shortening is a direct consequence of treatment-induced intracellular oxidative stress.

### Detection of persistant TBHP-induced oxidative modifications post fixation by *T***_1_** relaxometry

The transient nature of ROS makes the detection of short-lived radicals challenging in fixed-cell samples using conventional approaches. In our earlier experiments, where NDs were introduced into live cells prior to fixation, treatment with TMZ or TBHP consistently resulted in a decrease in the *T*_1_ relaxation time. Although this decrease is typically interpreted as increased magnetic noise arising from short-lived radicals in the live cells, it may instead reflect persistent intracellular modifications induced by ROS in the fixed cell samples.

We investigated whether timing of ND introduction influences the observed *T*_1_ response, by introducing NDs only after fixation(Fig.4a). To facilitate intracellular ND uptake in fixed cells, two delivery strategies were evaluated: direct incubation of NDs following fixation to allow passive diffusion, and membrane permeabilization using Triton X-100 to enhance intracellular entry. As shown in Fig.4c, *T*_1_ relaxometry measurements obtained using both delivery approaches yielded comparable results, indicating that the method of ND introduction post fixation does not significantly influence relaxation measurements in control cells. To confirm successful ND internalization under both conditions, fluorescence imaging was performed, demonstrating similar intracellular ND distribution across both conditions (Supplementary Fig.S6).

**Figure 4:**
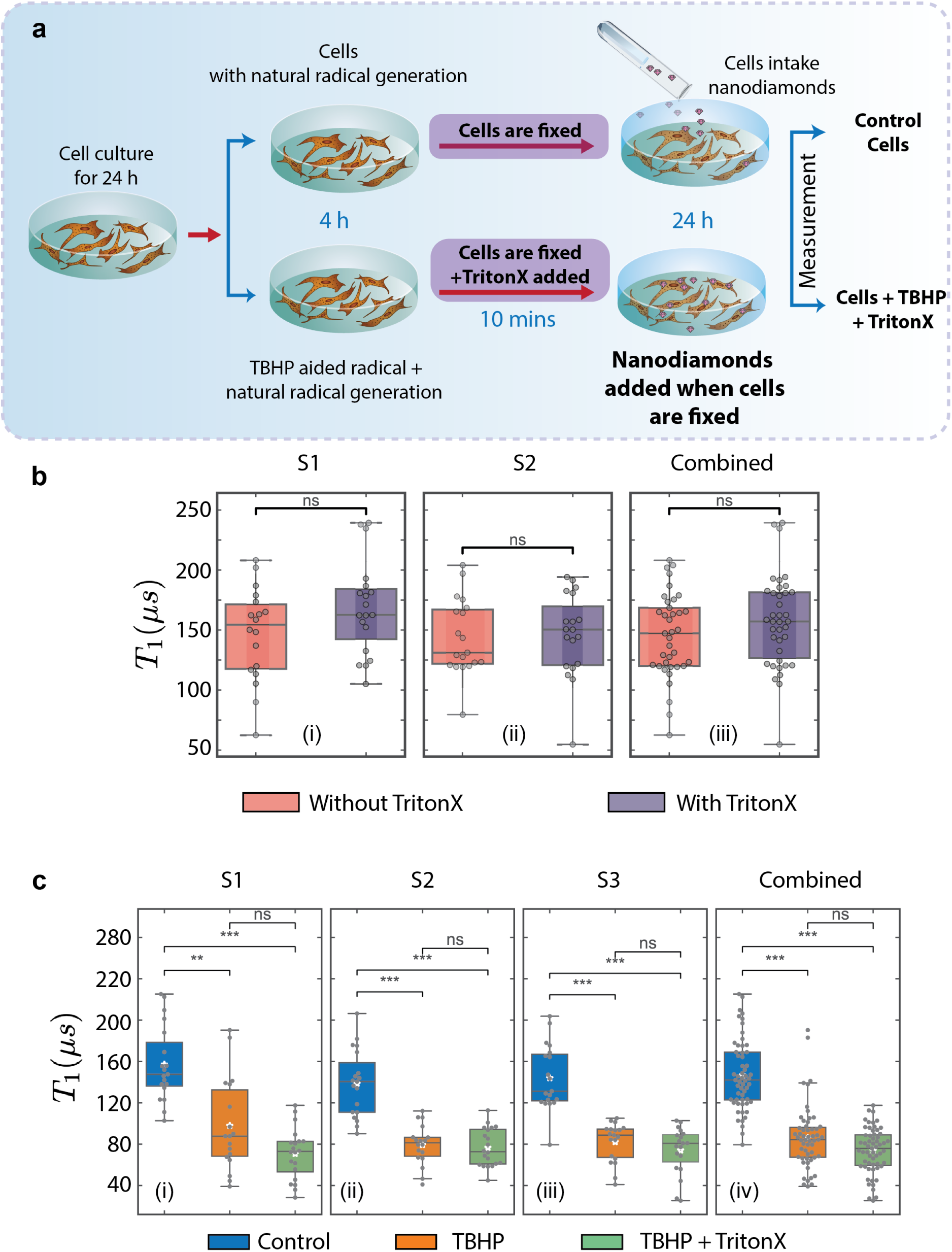
Detection of pesistant TBHP-induced oxidative modifications after fixation by. *T*_1_ **relaxometry**. (a) Schematic illustration of TBHP-induced radical generation and the experimental workflow for post-fixation sample preparation for relaxometry measurements. (b) *T*_1_ relaxometry measurements for U87-MG cells with and without treatment of TritonX across two independent experimental sets (S1 and S2) and the pooled dataset (Combined). Box plots show the distribution of *T*_1_ values (median), with individual measurements (gray dots) overlaid. (c) *T*_1_ relaxometry measurements for control and TBHP-treated U87- MG cells (with and without TritonX) across three independent experimental sets (S1–S3) and the pooled dataset (Combined). Box plots show the distribution of *T*_1_ values (median), with mean (white dot) and individual measurements (gray dots) overlaid. Colors indicate conditions as shown in the legend. For both (b) and (c): Statistical significance is indicated by adjusted *p*-values (Holm correction); thresholds: *∗ ∗ ∗ <* 0.001, *∗∗ <* 0.01, *ns >* 0.05. Colors indicate conditions as shown in the legend.

Based on these observations, subsequent comparative experiments were conducted in TBHP-treated samples, in which NDs were introduced after fixation using either passive diffusion or Triton X-100–mediated permeabilization, and the results were compared with control cells in which NDs were introduced post-fixation without Triton X-100. Under post-fixation delivery conditions, any observed change in *T*_1_ is expected to arise from persistent intracellular modifications rather than transient radical species. Consistent with this expectation, TBHP-treated samples post fixation also exhibited a clear reduction in *T*_1_ relaxation time relative to control (Fig.4c). Importantly, the cumulative trend across three independent replicates closely matched that observed when NDs were introduced prior to fixation.

Collectively, these results demonstrate that ND-based *T*_1_ relaxometry reliably detects long-lasting intracellular changes induced by oxidative stress. The consistent response observed across different ND delivery conditions supports the conclusion that this method captures persistent oxidative signatures rather than transient radical species. This capability provides a clear advantage over conventional ROS assays that depend on short-lived intermediates and highlights the potential of ND-based *T*_1_ relaxometry for evaluating drug response and oxidative stress at the single-cell level in fixed cellular systems, offering direct translational relevance for pathology-compatible analyses.

## Discussion and Conclusion

This study establishes, for the first time, NV-center–based *T*_1_ relaxometry using fluorescent NDs as a sensitive and reproducible platform for detecting chemotherapy-induced oxidative changes in GBM cells. TMZ treatment produced a clear dose-dependent reduction in *T*_1_ relaxation time in U87-MG cells, and this trend remained consistent across independent biological replicates despite the expected stochastic variability associated with single-particle intracellular measurements. Systematic optimization of ND dispersion, intracellular uptake, and photon-count normalization ensured that the observed changes in *T*_1_ reflect biologically driven alterations in the intracellular magnetic environment rather than measurement artifacts. Together, these findings demonstrate that ND-based relaxometry can provide a robust and quantitative readout of oxidative stress at the single-cell level.

A central advance of this work is the extension of quantum sensing beyond transient ROS measurements to the detection of persistent oxidative signatures that remain preserved after cellular fixation. Conventional ROS assays such as DCFDA remain widely used for monitoring intracellular oxidant production in live cells; however, their broad reactivity, susceptibility to auto- and photo-oxidation, dependence on intracellular esterases, and incompatibility with fixed samples limit their reliability and interpretability in many experimental and clinical settings.^51,52^ In contrast, *T*_1_ relaxometry detects paramagnetic species generated downstream of ROS activity, including stable redox-derived intermediates that persist beyond the lifetime of short-lived radicals. The persistence of *T*_1_ shortening following both TMZ and TBHP treatments, even when NDs were introduced after fixation, strongly supports the interpretation that the measured signal originates from long-lived intracellular oxidative modifications rather than transient ROS alone. The equivalence of results obtained using passive diffusion and Triton X–mediated permeabilization further confirms that the sensing response is independent of delivery strategy and reflects intrinsic cellular chemistry. These findings align with the broader biochemical understanding of TMZ-induced oxidative stress,^38^ which involves the formation of both transient radicals and stable lipid peroxidation products such as 4-HNE and MDA. Such aldehydic intermediates form covalent adducts with proteins and membranes and are increasingly recognized as major contributors to treatment-induced cytotoxicity. Importantly, detoxification of these reactive aldehydes by enzymes such as ALDH1A3 has been strongly linked to TMZ resistance and tumor relapse in GBM.^53^ In this framework, the detection of persistent oxidative footprints through ND-based relaxometry provides a mechanistically relevant measure of chemotherapy-induced damage. Conversely, the absence or attenuation of *T*_1_ shortening may reflect enhanced antioxidant defenses characteristic of resistant cell populations. Thus, beyond detecting oxidative stress, this platform offers the potential to function as an early indicator of therapeutic resistance and treatment outcome.

Beyond its immediate application to GBM, this platform introduces a broader paradigm for retrospective redox analysis in fixed biological specimens. The ability to detect stable oxidative signatures in fixed cells offers clear advantages for compatibility with pathology workflows, including archival specimen analysis and longitudinal clinical studies. Because many chemotherapeutic strategies rely on oxidative mechanisms, the approach described here may be adaptable to other cancer types and therapeutic contexts where redox imbalance plays a critical role. Furthermore, the nanoscale sensitivity of NV centers provides a unique opportunity to interrogate cellular heterogeneity, an increasingly recognized determinant of treatment response and disease progression. Future studies may expand this framework in several directions. Functionalization of nanodiamonds with organelle-specific targeting motifs could enable spatially resolved mapping of oxidative stress within defined intracellular compartments. Integration with live-cell imaging modalities may further allow dynamic monitoring of redox fluctuations during treatment cycles. In addition, extending this methodology to patient-derived cells or tissue biopsies could establish its clinical relevance for personalized therapeutic decision-making. Ultimately, the convergence of quantum sensing technologies with cellular and clinical biology has the potential to redefine how oxidative processes are measured, interpreted, and translated into actionable biomedical insights.

## Methods

### Drop-casting nanodiamonds on coverslips

70 nm nanodiamonds from Sigma-Aldrich were drop-casted between two 22 *mm*^2^ coverslips. For drug interaction with media studies, a drop of 70 *µl* consisting of properly mixed 10 *µg/ml* NDs and 250 *µM/*1000*µM* TMZ was cast between two 22 mm² coverlips along with mounting media. The sample was then mounted on the confocal microscope, and *T*_1_ measurements were performed as described in the following section.

### Cell culture

U87-MG cells, an aggressive Grade IV glioblastoma cell line, were kindly provided by Prof. Shilpee Dutt’s lab(ACTREC, Navi Mumbai). The cells were cultured in DMEM (HiMedia) supplemented with 10% fetal bovine serum (HiMedia), 10% antibiotic-antimycotic solution (Gibco), and 1% L-glutamine (Gibco). They were maintained at 37*^◦^*C in a humidified incubator with 5% *CO*_2_. For experiments, cells were seeded onto 22*mm*^2^ plastic coverslips at a density of 60,000 cells per coverslip. Depending on the experiment, cells were harvested at various time points following treatment with either NDs or Temozolomide TMZ.

### ND treatment and processing

U87-MG cells were seeded on 22 mm² coverslips and allowed to adhere for 24 hours, as described previously. For internalization studies, 35 nm and 70 nm nanodiamonds (NDs; Sigma-Aldrich) were initially treated at a concentration of 10 µg/mL. The NDs were diluted from a 1 mg/mL stock (provided by Sigma) to a working concentration of 400 µg/mL in fetal bovine serum (FBS). This working solution was subjected to sonication for 1 hour, followed by vortexing for 30 minutes to disperse aggregates before cell treatment. However, upon imaging, significant aggregation was observed using this approach. To address this, an optimized protocol was developed. The ND-FBS working solution was first sonicated for 1 hour, then vortexed for 1–2 minutes, and subsequently left undisturbed at room temperature. Without any further mixing, aliquots were carefully collected from the supernatant after 0, 5, 10, and 30 minutes of settling. Among these, the 30-minute timepoint yielded the best dispersion results, as detailed in the Results section. Based on these observations, all subsequent treatments were performed using the 30-minute supernatant and at a reduced ND concentration of 5 µg/mL to further minimize aggregation and improve cellular uptake. NDs were incubated for 2 hours post-treatment (unless mentioned otherwise). Post 2 hours of incubation, the cells were fixed with 4% formaldehyde. After fixation, the cells were washed twice with 1× PBS and then stained with Hoechst (Sigma). The coverslips were then mounted on glass slides for imaging. Imaging was performed at 20X in the Eclipse Tie2 Nikon and at 60X magnification in the Leica confocal and PicoQuant time-resolved FLIM microscopes.

### MTT assay

To assess the viability of U87-MG cells following nanodiamond (ND) treatment, an MTT (dimethyl thiazol diphenyl tetrazolium bromide) assay was carried out. Approximately 5,000 cells were seeded per well in a 96-well plate. After allowing the cells to adhere and grow for 24 hours, they were treated with varying concentrations of NDs (ranging from 0 to 10 *µ*g/mL). Following ND exposure, cells were incubated for another 24 hours. After the treatment period, MTT reagent (Sigma) was added to each well at a final concentration of 0.5 mg/mL, and the plates were incubated for 4 hours at 37°C. Formation of purple formazan crystals was confirmed under a microscope. The media containing the MTT reagent were then carefully removed, and 200 *µ*L of DMSO was added to each well to dissolve the crystals. Absorbance was read at 570 nm using a SpectraMax M5 (Molecular Devices company) plate reader, and the percentage of viable cells was calculated based on the absorbance values.

### Fluorescent imaging and in-cellulo analysis of NDs

To prepare the sample, 70 nm nanodiamonds were introduced into the cells at a concentration of 10 *µ*g/ml, 24 hours post-seeding. 2 hours post-treatment, cells were fixed and processed for imaging. During processing, Hoechst dye was added at a 1:3000 ratio, which stains the nucleus and is subsequently mounted on a slide with antifade. The sample was viewed under the PicoQuant MicroTime 200 confocal microscope, using a 404/532 double dichroic mirror along with a 425 Long Pass filter for Hoechst dye imaging and a 532 Long Pass filter for nanodiamonds. 2 regions were selected from 2 slides, and each was imaged twice; once for the nucleus and once for the nanodiamonds. Images include 1 to 5 cells and information about the counts and lifetime. Images are taken at the following timepoints after fixation: 8, 26, 50, and 74 hours. This process was repeated for two sets.

The intensity and area of NDs in-cellulo were analyzed using ImageJ. Cells were identified by the fluorescent images of the Hoechst-stained nucleus. Nanodiamonds within each cell were then selected by applying a threshold to the photon counts, converting the image to a binary format where pixels above the threshold represented NDs, and defining them as Regions of Interest (ROIs). The threshold was chosen as 17.33*±*2.79 and 18.53*±*1.11 counts for sets 1 and 2, respectively, after calibrating counts with grayscale values. The scale was set in ImageJ using the scale bar in the image, allowing accurate calculation of the physical area occupied by NDs within each ROI. These ROIs were then overlaid onto the original fluorescence image to measure the average fluorescence intensity (counts) of nanodiamonds in each cell. The resulting data, including ND area and intensity values, were exported and further analyzed using Python.

### Temozolomide and TBHP treatments

As described earlier, U87-MG cells were seeded onto 22 mm² coverslips at a density of 60,000 cells per coverslip. After 24 hours of incubation, cells were treated with TMZ at concentrations of 0 µM (control), 250 µM, and 1000 µM. TBHP (positive control for ROS generation) was treated at 0 µM (control) and at 100 µM. Following 24 hours of TMZ treatment and 4 hours of TBHP treatment, the cells were fixed using 4% formaldehyde. Nanodiamonds (NDs) were added 2 hours prior to fixation at a concentration of 5 µg/mL. After fixation, the cells were washed twice with 1× PBS, and the coverslips were mounted using Fluoromount Aqueous Solution (Sigma) onto fresh 22 mm² coverslips. These samples were then used for *T*_1_ relaxometry measurements.

### ND uptake post fixation of cells

To check the cellular uptake of NDs post fixation, the cells were seeded for 24 hours on 22 mm² coverslips, after which they were fixed for 10 minutes with 4% Formaldehyde following a thorough 1X PBS wash, thrice, each for 5 minutes. Post fixation, we administered NDs to one set, and another set was first permeabilized with 0.1% Triton X-100 for 10 minutes, followed by similar washing steps with 1X PBS, and then NDs were administered. NDs were provided to the cells at a concentration of 10 *µg/ml* in complete media for 24 hours. We also performed similar experiments with TBHP treatment, which was provided for 4 hours, post 24 hours of cell seeding, after which similar fixation and permeabilization steps were performed. Post ND incubation, the cells were washed twice with 1X PBS and then stained with Hoechst (Sigma). Coverslips were mounted on slides with Fluoromount and were used for imaging and *T*_1_ measurement.

### ***T*_1_** relaxometry experiment and analysis

The relaxometry experiments with NV centers in diamond were performed using a custom-built confocal setup (Supplementary Fig.S7). The data was collected by custom-written Python scripts interfaced with a National Instruments data acquisition card (NI-DAQ). For each run, an automated pulse sequence was applied to the Acousto-Optic Modulator (AOM) to modulate the laser. The pulse sequence applied consists of optical initialization, variable-duration dark times (logarithmically spaced between 300ns and 3ms), and subsequent optical readout. The typical measurement consists of 3 runs with 25 timesteps and 40000 averages per step per run. The resulting photon counts were used to calculate the contrast according to the formula: 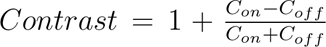 where *C_on_* and *C_off_* correspond to the photon counts with and without optical initialization, respectively. Simultaneously, the brightest spot in a 1 *µ*m^2^ region was tracked using Qudi’s tracking algorithm every 15 seconds to ensure data were collected from the same location and to compensate for mechanical drift. The data were saved in the ‘hdf5’ format, including raw counts (for *C_on_* and *C_off_*), time step, and contrast. The contrast decay curve was fitted with the single exponential decay function; *C*(*τ*) = *a* exp (*−τ/T*_1_) + *b*. Where *a* and *b* are fit parameters and *τ* is the dark time. The SciPy curve-fitting routine was used to fit the curve and extract the *T*_1_ relaxation time. To ensure data quality, the coefficient of determination (*R*^2^) was calculated for each fit, and only measurements with *R*^2^ *>* 0.8 were included in the analysis. Data visualization and the box plots were generated using custom Python scripts.

### DCFDA assay

U87-MG cells were seeded into 96-well plates at densities of 10,000 cells per well for TBHP and TMZ treatment, allowing them to adhere for 24 hours. Treatments were then carried out as described earlier. After 24 hours of TMZ treatment and 4 hours of TBHP treatment, the DCFDA assay was performed following the manufacturer’s protocol (Abcam). Briefly, cells were first incubated with 1× DCFDA buffer for 20 minutes to equilibrate. After removing the buffer, DCFDA was added at a final concentration of 20 µM, freshly prepared in 1× DCFDA buffer. The plate was incubated in the dark at 37°C for 45 minutes. Following incubation, the DCFDA solution was removed, and cells were washed and overlaid with 1× supplemented buffer (DCFDA buffer containing 10% FBS). Fluorescence was then measured in endpoint mode at an excitation/emission wavelength of 485/535 nm using a SpectraMax M5 (Molecular Devices) plate reader.

## Acknowledgement

We deeply acknowledge Prof. Shilpee Dutt (ACTREC Navi Mumbai/ JNU, New Delhi), who kindly helped us by providing the GBM cell line U87-MG for this study. We acknowledge Alexander Franzen for images of optical components from the Component Library. We acknowledge our project student Avni Kulkarni for her help during the initial experimental standardizations. KS, ST, and AM acknowledge funding from Wadhwani Research Center for Bioengineering. In addition, KS acknowledges funding from DST National Quantum Mission and AOARD grant number FA2386-23-1-4012. AM thanks the Science and Engineering Research Board (ANRF-SERB) for research funding (Grant No. CRG/2021/009001-II).

## Author contributions

JP and AG contributed equally to this work. They planned and conducted the experiments and analyzed the data. SS contributed to the biological experiments. Ay.M and DZ collected the FLIM images. DZ also contributed to the initial characterization data. JP and AG wrote the manuscript with inputs and edits from all coauthors. AM, ST, and KS conceptualized the idea. All authors reviewed the manuscript. AM supervised bio related aspects of the work. KS supervised all aspects of the work.

## Additional information

Supplementary information available.

