## Supplementary Information for "Quantum Sensing Reveals Chemotherapy Induced Persistent Oxidative Footprints in Fixed Brain Cancer Cells"

### Supporting Information Available

#### Supplementary Figure 1

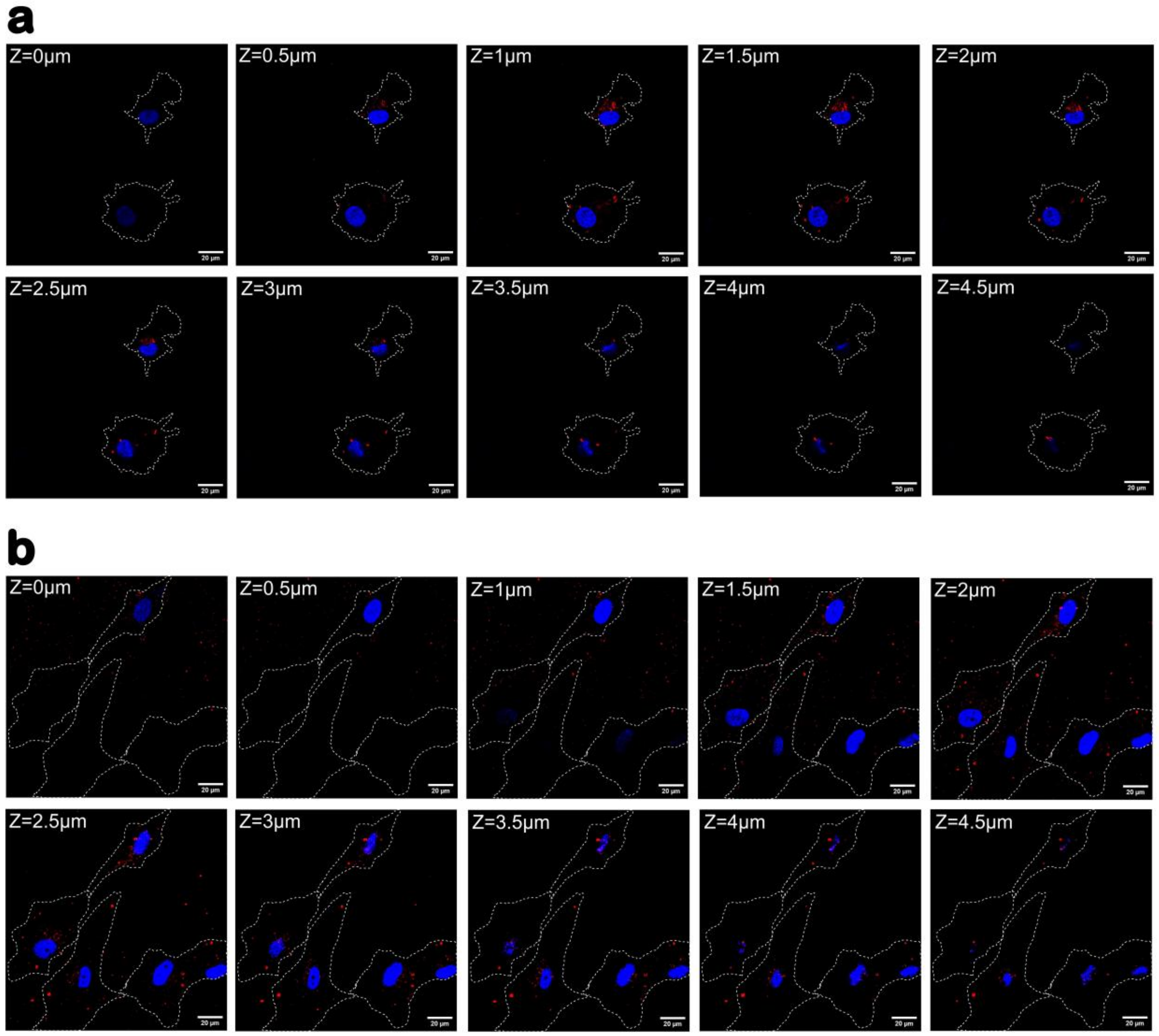


Figure S1: **Cellular Uptake of Nanodiamonds in U87-MG GBM Cells.** (a, b) 70nm NDs at 10 $\mu$g/ml. Z-stacked images reveal the presence of NDs within cells. Red fluorescence corresponds to the ND signal, blue fluorescence corresponds to the Hoechst-stained nucleus, and the cell boundary is represented by white dashed lines, adapted from DIC images. Scale bar corresponds to 20 $\mu$m.

#### Supplementary Figure 2

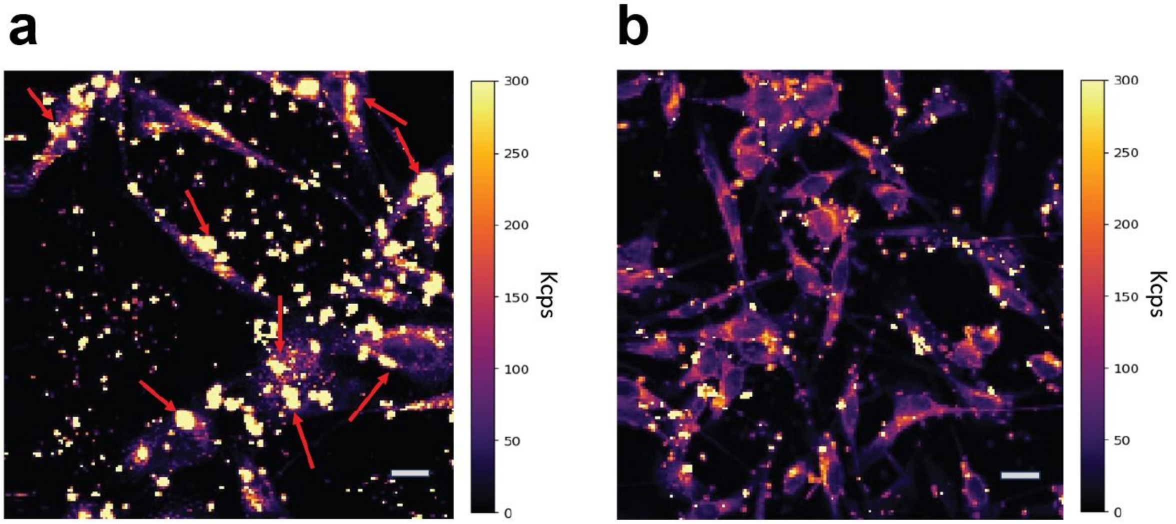


Figure S2: **Intracellular characterization of NDs in U87-MG cells for reliable T1 relaxation measurements.** (a) The confocal scan (scale bar 20$\mu m$) of cells containing nanodiamonds (NDs). The term ”Kcps” refers to kilo counts per second. Relatively large clusters of NDs are clearly observed within the cells, as indicated by the red arrow. (b) The confocal scan (scale bar 20$\mu m$) displays a scarcity of clusters, predominantly located outside the cells. This is achieved by following the steps mentioned in the ND treatment and processing section.

#### Supplementary Figure 3

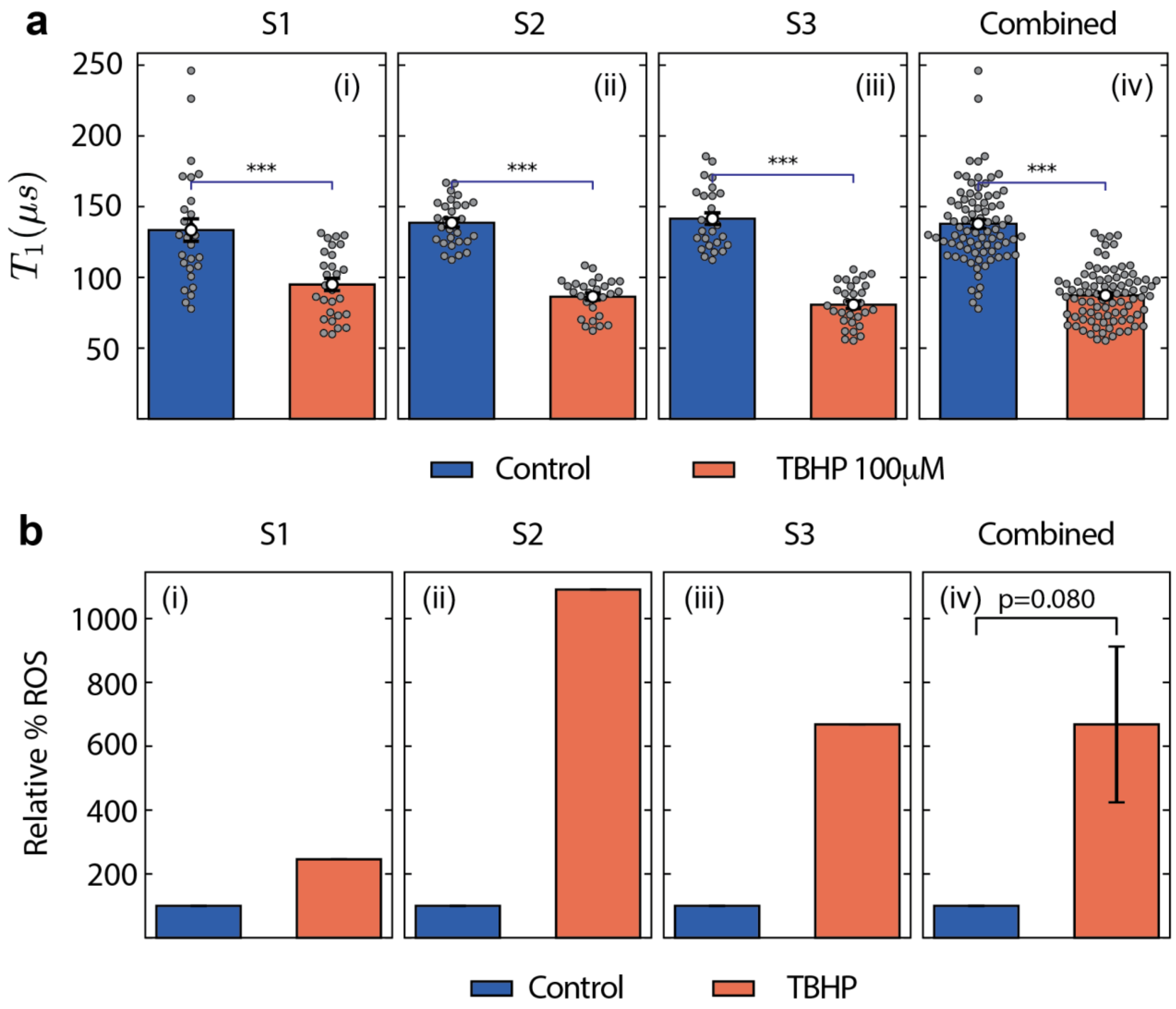


Figure S3: **T1 relaxometry and DCFDA measurements from positive control TBHP.** (a) $T_1$ relaxometry measurements for control and TBHP-treated U87-MG cells (100 $\mu$M) across three independent experimental sets (S1–S3) and the pooled dataset. A significant reduction in $T_1$ is observed upon TBHP treatment ($***p < 0.001$). Bars represent mean (white dot) ± SEM, with individual measurements shown as gray dots. (b) DCFDA assay showing intracellular ROS levels for control and TBHP-treated cells (100 $\mu$M), including individual biological replicates (S1-S3) and the combined analysis. Error bar represents ± SEM. $p - value = 0.08$

#### Supplementary Figure 4

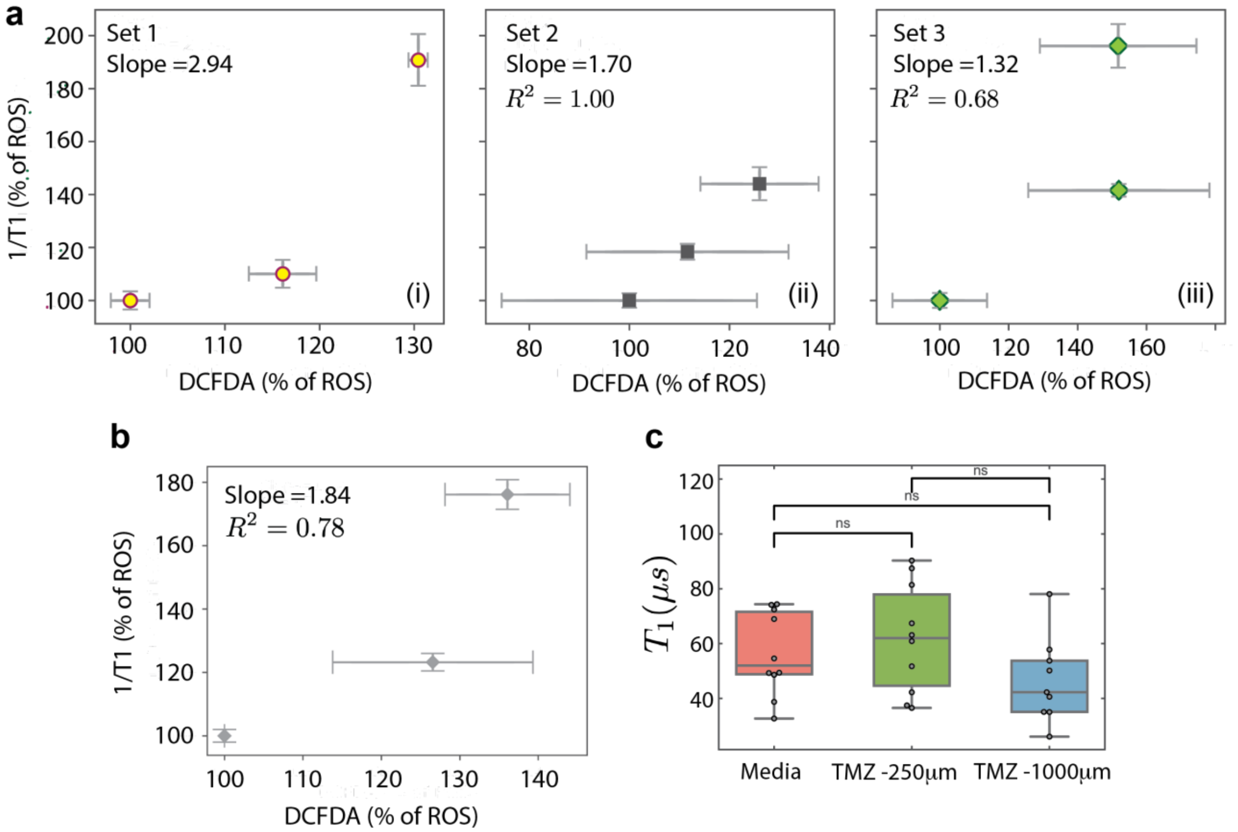


Figure S4: **Comparison of $T_1$ relaxometry and DCFDA with $T_1$ in TMZ as negative control** (a) Correlation between DCFDA assay (% ROS) and normalized inverse relaxation time ($1/T_1$, expressed as % ROS) across three independent experimental sets (Set 1–3), shown individually. (b) Combined analysis of all three datasets, showing the overall correlation between DCFDA and $T_1$-based measurements. For both (a) and (b): Data points represent mean ± SE; corresponding slopes and $R^2$ values are indicated. Regression lines are omitted for clarity. (c) $T_1$ relaxometry measurements in cell-free medium containing nanodiamonds with TMZ (0, 250, and 1000 $\mu$M), serving as the negative control. Box plots show median and distribution, with individual measurements overlaid (gray dots). *ns*, not significant ($p > 0.05$).

#### Supplementary Figure 5

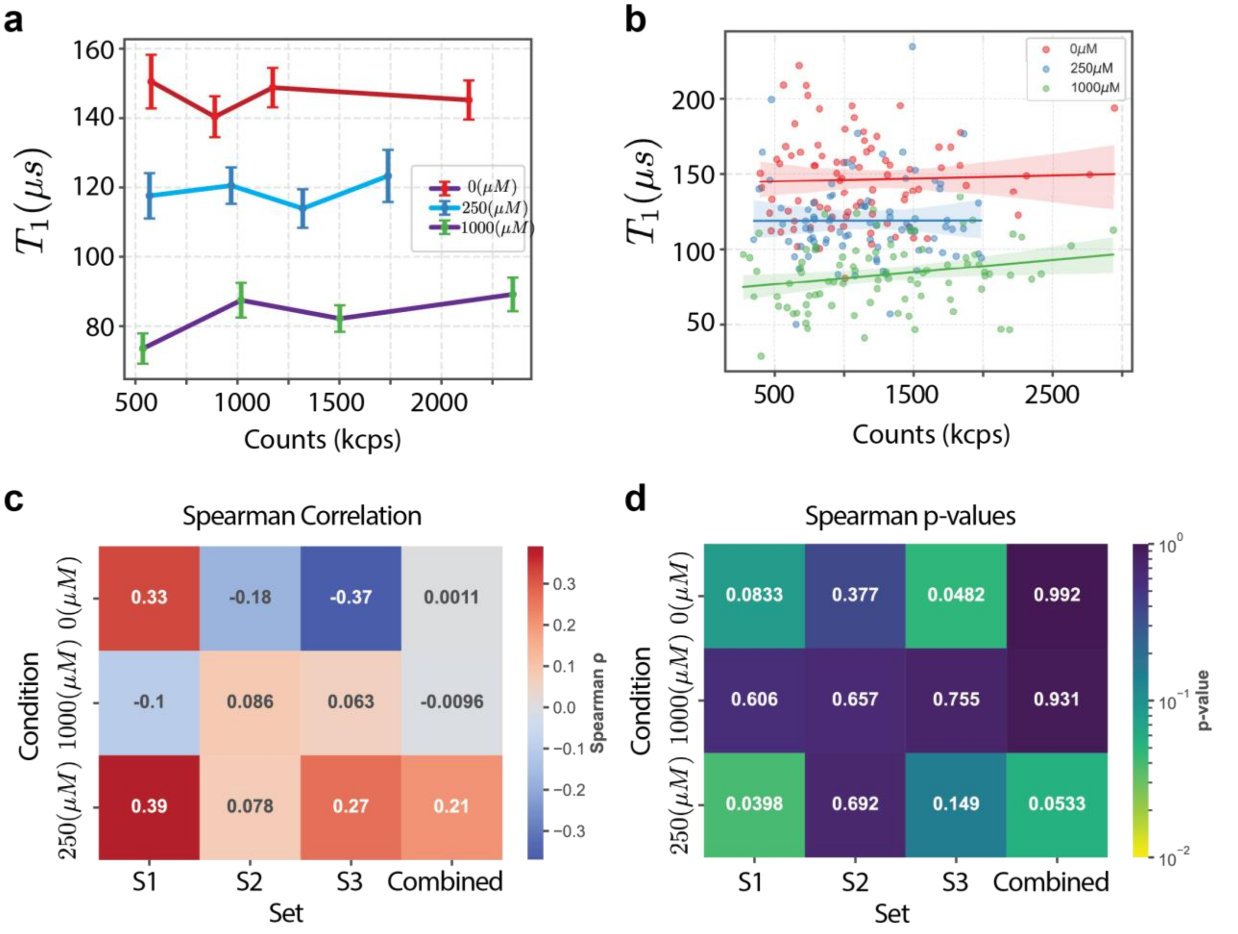


Figure S5: **Assessment of fluorescence (intensity) effects on $T_1$ relaxometry.** (a) Binned analysis of $T_1$ as a function of photon counts (kcps) for the pooled dataset (S1–S3), grouped into quartiles and shown as mean ± SEM for each condition (0, 250, and 1000 $\mu$M). No systematic variation in $T_1$ is observed across intensity bins. (b) Scatter plots of $T_1$ versus photon counts with linear regression fits and 95% confidence intervals, demonstrating the absence of any monotonic dependence across all conditions. (c) Spearman correlation coefficients ($\rho$) for individual datasets and the combined data reveal weak, inconsistent correlations without a reproducible trend. (d) Corresponding Spearman $p$-values show occasional significance in isolated cases; however, neither the pooled dataset nor the overall trends reach statistical significance, confirming that fluorescence intensity does not bias $T_1$ measurements.

#### Supplementary Figure 6

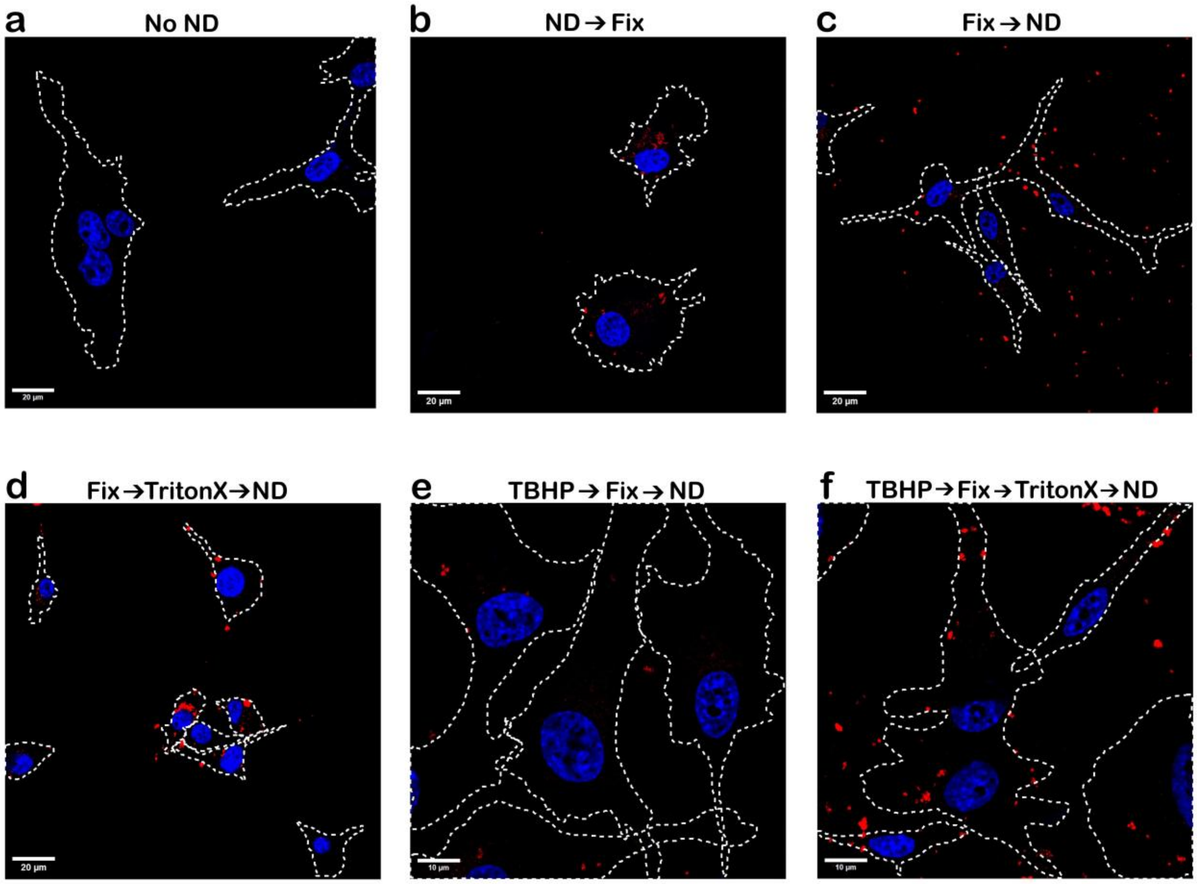


Figure S6: **Effect of fixation and permeabilization on nanodiamond uptake in U87MG cells.** (a) Negative control (no NDs). NDs were introduced (b) prior to fixation, (c) post-fixation, and (d) post-fixation with TritonX permeabilization. TBHP-treated cells with NDs added (e) post-fixation and (f) post-fixation with permeabilization. Pseudo-colors: blue corresponds to Hoechst-stained nucleus; red corresponds to NDs. White dashed lines denote cell boundaries extracted from DIC images. Scale bars: (a–d), 20 $\mu$m; (e,f), 10 $\mu$m.

#### Supplementary Figure 7

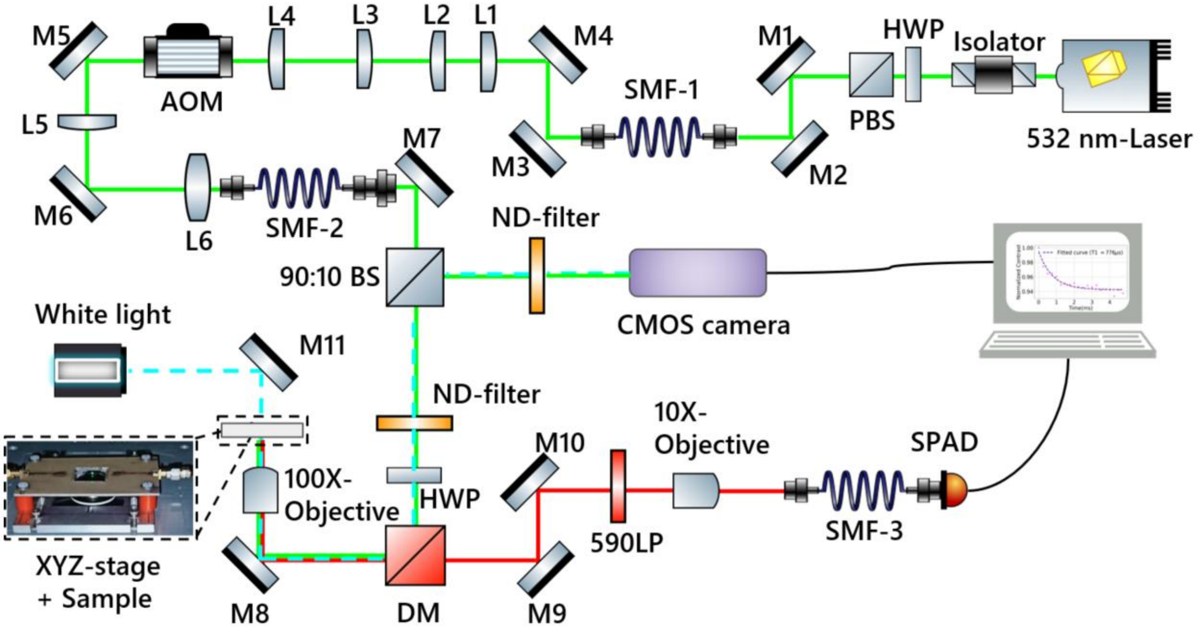


Figure S7: **Schematic for custom-built confocal setup.** The green path corresponds to the 532 nm laser used to excite the sample, and the emission path is shown by the red line. The path for white light is shown by the cyan dashed line. Abbreviations: HWP — half-wave plate; PBS — polarizing beam splitter; M — mirror; SMF — single-mode fiber; L — lens; AOM — acousto-optic modulator; BS — beam splitter; ND filter — neutral density filter; DM — dichroic mirror; LP — long-pass filter; SPAD — single-photon avalanche diode.
